# Atypical cortical encoding of speech identifies children with Dyslexia versus Developmental Language Disorder

**DOI:** 10.1101/2022.10.26.513864

**Authors:** João Araújo, Benjamin D Simons, Varghese Peter, Kanad Mandke, Marina Kalashnikova, Annabel Macfarlane, Fiona Gabrielczyk, Angela Wilson, Giovanni M. Di Liberto, Denis Burnham, Usha Goswami

## Abstract

Slow cortical oscillations play a crucial role in processing the speech envelope, which is perceived atypically by children with Developmental Language Disorder (DLD) and developmental dyslexia. Here we use electroencephalography (EEG) and natural speech listening paradigms to identify neural processing patterns that characterize dyslexic versus DLD children. Using a story listening paradigm, we show that atypical power dynamics and phase-amplitude coupling between delta and theta oscillations characterize dyslexic and DLD children groups, respectively. We further identify EEG common spatial patterns (CSP) during speech listening across delta, theta and beta oscillations describing dyslexic versus DLD children. A linear classifier using four delta-band CSP variables predicted dyslexia status (0.77 AUC). Crucially, these spatial patterns also identified children with dyslexia in a rhythmic syllable task EEG, suggesting a core developmental deficit in neural processing of speech rhythm. These findings suggest that there are distinct atypical neurocognitive mechanisms underlying dyslexia and DLD.

## Introduction

Developmental Dyslexia and Developmental Language Disorder (DLD) affect between 3-7% of the population, negatively impacting education and life chances [1, 2]. Behavioural research shows that atypical linguistic processing lies at the heart of both disorders. While dyslexia is diagnosed primarily on the basis of difficulties in reading and spelling once schooling begins, at-risk children show atypical processing of phonology [3, 4] (the sound structure of speech) from infancy [5, 6]. By contrast, DLD children show difficulties in acquiring oral language from infancy, particularly grammar: they are often “late talkers” [7]. Crucially, these difficulties cannot be attributed to lower intellectual ability, inadequate schooling, or overt hearing impairment. However, both disorders are characterized by impairments in processing the speech amplitude envelope, particularly amplitude envelope rise times, and associated difficulties in processing speech rhythm [8, 9].

For both dyslexia and DLD, it has been hypothesized that these impairments are related to atypical neural oscillatory responses to the speech envelope for children: ‘Temporal Sampling’ (TS) theory [10–12]. Children with both disorders show impairments in perceiving syllable stress [13], a fundamental component of speech rhythm carried by low-frequency amplitude modulations (AMs) in the speech envelope. Prior research with dyslexic children also shows atypical cortical tracking of low-frequency speech envelope information across languages [14–19]. Such phenomenology may be related to impaired neural mechanisms of speech edge detection found in dyslexia [20]. Atypical neural processing of speech is thought to result in the development of speech-based representations that encode poorly syllable stress patterns (dyslexia) and/or prosodic phrasing (DLD) [21, 22].

Previous electrophysiology studies with DLD children have shown atypical event-related potential responses as well as reduced sensitivity to temporal asynchronies in audio and visual scenes, especially at longer timescales [23]. However, to date, no studies have pinpointed neural mechanisms capable of distinguishing children with dyslexia from children with DLD based on temporal parameters in naturalistic speech conditions. To this end, we examined the relative magnitude and coupling of low frequency neural oscillations – namely delta and theta, those identified by TS theory – during receptive speech tasks. Different temporal rates are thought to support parsing of the speech signal into linguistic units; for example, stressed syllables are in the range 1-4Hz, syllables 4-8Hz, and phonemes >30Hz [24–26]. While adult studies find an increase in neural oscillatory power at similar frequencies to segregate speech information in relevant time windows (delta in the range 1-4Hz, theta 4-8Hz, and gamma >30Hz) [27], recent work highlights the relative increase in the magnitude of delta (and not theta) oscillations when processing meaningful (but not unintelligible) speech, suggestive of a role in top-down syntactic processing [28]. Furthermore, studies of auditory attention (i.e., cocktail party paradigms) show that delta and theta oscillatory responses are modulated by selective attention processes [29, 30]. Oscillatory cross-frequency coupling (CFC) is also hypothesized to play a role in decoding the speech signal at a more sensory (bottom-up) level [27, 31, 32]. CFC has a role in sensory stimulus selection, indirectly regulating neural excitability by controlling the magnitude of oscillations at different timescales [33]. Neurally, the primary auditory cortex is organized in phase-amplitude hierarchies, so that the delta phase modulates theta amplitude and the theta phase controls gamma amplitude [31]. Changes in low-frequency oscillatory coupling (namely delta and theta) have both been reported when stimulus rhythm is relevant to the task [33] and in long time windows of audio-visual stimuli integration [34]. Therefore, studying this phenomenology in both DLD and dyslexia is likely to shed light on the nature of the sensory-neural deficits underlying both disorders.

Electrophysiology studies with dyslexic children may also provide useful biomarkers for accurate diagnosis, similarly to studies conducted with Autism Spectrum Disorder [35] or Parkinson’s disease [36]. In fact, recent work has shown that it is possible to build dyslexia classifiers from the EEG of dyslexic children listening to rhythmic auditory stimuli (white noise) presented at different rates [37, 38]. These studies use several different types of features (time, frequency, fractal or CFC graph networks), with large and complex non-linear models, achieving good performance metrics. However, due to the nature of these approaches, the connection between model parameters for neural data and the underlying cognitive/linguistic processes is not always transparent.

Here, we sought to explore how relationships between oscillatory rhythms are affected in both dyslexia and DLD. We aimed to develop an interpretable EEG classifier that can distinguish reliably between children with dyslexia and typically-developing children. By comparing spatial patterns of oscillatory activity from different receptive speech tasks, we also aimed to shed light on how the estimated biomarkers might relate to cognitive/linguistic processes. Specifically, we compared the neural oscillatory responses hypothesized as core by TS theory of children with dyslexia versus DLD in natural speech tasks. We predicted that the relative amplification of theta versus delta responses and delta-theta cross-frequency coupling might be altered in dyslexia and DLD. Our findings show that distinct dynamics between slow oscillatory rhythms selectively segregate dyslexic (theta/delta power) and DLD children (delta-theta phaseamplitude coupling) from typically-developing children in a story listening task at the group level. We next hypothesized that delta and theta oscillatory responses to connected speech might provide sufficient information to classify neural patterns that identify dyslexia at the single trial level. Consistently, we found that dyslexic and typically-developing children can be identified based on oscillatory responses for connected speech across these bands using a simple linear classifier. Finally, we explored how specific or universal these discriminatory patterns might be to natural language versus general speech rhythm processing. Here, we used EEG from a rhythmic syllable repetition task collected with new dyslexic and typically-developing children. By training a classifier on the EEG from rhythmic syllable listening using discriminatory features derived from story listening, we find a generalization of features for classification. In addition to providing a general classifier to identify dyslexia, our modelling suggests that classification of dyslexia is related to atypical neural processing of rhythm (rather than semantic or syntactic processing).

## Results

### Dyslexic but not DLD children show different theta/delta oscillatory power ratios during speech processing

Our first goal was to understand how the relationships between low-frequency oscillations during speech processing change with developmental dyslexia or DLD. To this end, we analysed EEG data collected as DLD, dyslexic and typically-developing children listened to a story (Figure 1a, see STAR Methods for more details). In line with previous studies in adults and children [15, 28], we focused on delta (1-4Hz) and theta (4.5-8Hz) power dynamics, exploring differences between typically-developing children, children with dyslexia and children with DLD. Age and neuropsychological profiles for our groups are shown in Table 1 (see Table S1 for the full behavioural dataset). To focus the analysis, we placed emphasis on the uncorrelated brain regions that provided the highest differential response to the listening task. To identify these regions, we made use of <u>P</u>rincipal Component Analysis (PCA) and computed band power from each PC individually. We retained only PC filter vectors with the highest eigenvalues, setting the threshold at a total of 70% variance explained (see STAR Methods for further details). Applied to the current dataset, this translated to just three PCs (Figure 1b). Their associated channel weights showed stereotypic patterns consistent with the audio-visual nature of the experimental task: PC1 (total variance explained: 50%) showed an enrichment of channel weights in the central region of the scalp, where evoked auditory potentials are usually observed. PC2 (total variance explained: 15%) showed a pattern of bilateral temporal electrodes covering both auditory cortices. Finally, PC3 (total variance explained: 13%) lay mostly on occipital channels.

**Figure 1.**
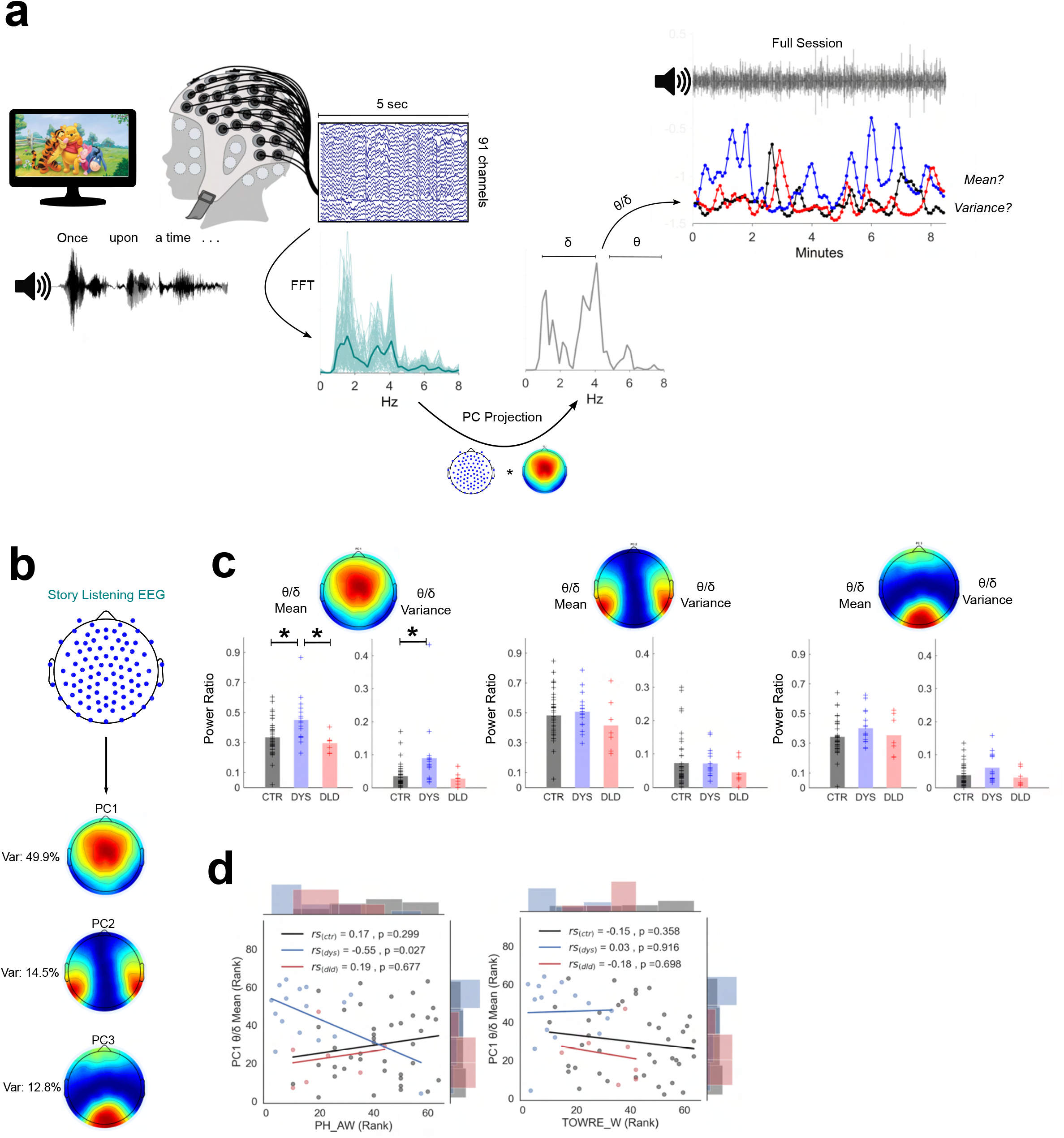
Dyslexic children show atypical theta/delta EEG power ratio during speech listening. (a) Story listening paradigm (Winnie-the-Pooh) and pseudo-online analysis pipeline (see STAR Methods). Dimension reduction analysis was applied to EEG data using principal component analysis. EEG projection to principal component (PC) 1 is used for illustration. Examples of story sessions from a dyslexic (blue), DLD (red) and a typically-developing child (black) are shown at top right. (b) Map of channel weights of the three most important principal components: weights and respective percentage of variance explained. (c) Theta/delta ratio mean and variance across all three PCs. Atypically high mean and variance is present for dyslexics on PC1 (CTR – typically-developing children, DYS – dyslexic children, DLD – DLD children). * p < 0.05 (Bonferroni-corrected). (d) Spearman correlations and uncorrected p-values between PC1 theta/delta and phonological awareness (leftmost panel) and reading (timed single word reading test, right panel).

**Table 1.** Story listening task: participant sample characterization.

|  | CTR |  | DYS | DLD |
| --- | --- | --- | --- | --- |
|  | RL<br>n = 13 (7F) | CA<br>n = 27 (10F) | n = 16 (6F) | n = 7 (2F) |
| **Age (months)** | 83.7 (6.0) | 113.6 (13.1) | 114.1 (18.5) | 95.0 (15.3) |
| **PH_AW<sup>a</sup>** | 106.4 (10.0) | 103.7 (10.5) | 86.0 (11.4) | 91.9 (8.0) |
| **TOWRE_W<sup>b</sup>** | 100.9 (15.6) | 102.6 (14.2) | 77.3 (11.1) | 94.4 (6.9) |
| **TOWRE_NW<sup>c</sup>** | 99.2 (14.7) | 102.5 (13.3) | 78.2 (8.2) | 94.7 (12.7) |
| **TROG<sup>d</sup>** | 106.5 (6.7) | 106.0 (8.6) | 101.1 (9.5) | 81.7 (16.1) |
| **WIAT_VOCAB<sup>e</sup>** | 107.5 (11.8) | 106.7 (11.3) | 102.4 (10.8) | 90.1 (12.4) |
| **CELF_SENTENCES<sup>f</sup>** | 11.2 (2.7) | 11.8 (2.3) | 9.4 (2.8) | 7.3 (2.3) |
| **NVIQ<sup>g</sup>** | 111.6 (11.4) | 114.4 (9.0) | 104.9 (9.3) | 99.0 (12.2) |

CTR – Typically Developing (control group); RL – Reading Level; CA – Chronological Age; DYS – Dyslexic children; DLD – Developmental Language Disorder. Standard deviations in parenthesis
<sup>a</sup> Comprehensive Test of Phonological Processing
<sup>b</sup> Test of Word Reading Efficiency: Words
<sup>c</sup> Test of Word Reading Efficiency: Non-Words
<sup>d</sup> Test for Reception of Grammar
<sup>e</sup> WIAT Vocabulary
<sup>f</sup> Clinical Evaluation of Language Fundamentals- Recalling Sentences (M = 10, SD = 3)
<sup>g</sup> Non-verbal IQ: Kaufman Brief Intelligence Test-Matrices

Theta/delta power ratios were calculated to test for the relative magnitude of theta oscillations compared to delta oscillations across groups. As many ratio distributions did not show a normal distribution (based on Shapiro-Wilk tests), nonparametric Kruskal-Wallis ANOVAs and post-hoc Wilcoxon tests were used to assess group differences. Two metrics were evaluated for each child for PC power ratios across the experimental session: the mean ratio value, to check for a general difference in the band power relationship trend; and the variance of this ratio, to test for differences in consistency for this relationship. The results are shown in Figure 1c. For PC1, group differences were found on the mean theta/delta ratio across groups (H = 10.96, p = 0.0042). Post-hoc Wilcoxon tests showed a higher mean theta/delta ratio for dyslexic children when compared to typically-developing children (Z = −2.87, p = 0.004) and DLD groups (Z = 2.77, p = 0.0056). Importantly, these differences did not arise because of a general difference in delta (H = 2.68, p = 0.26) or theta power (H = 1.39, p = 0.5). Group differences were also found regarding the consistency across epochs for theta/delta ratio variance (H = 11.49, p = 0.0032). Overall, ratio variances were higher for dyslexic children compared to typically-developing children (Z = −3.26, p = 0.0011). Differences in ratio variance between dyslexics and DLDs were not significantly different following Bonferroni correction (Z = 2.24, p = 0.0252). DLDs did not differ from typically-developing children regarding the power ratio consistency and mean value for PC1 (p > 0.05 for both), and no group differences were found for the average value or consistency of any other PC filters (p > 0.05). Taken together, these results suggest an atypically high and less consistent power dynamic between delta and theta oscillations for dyslexic children (but not for DLD) and that these differences are present in centrally-located regions of the scalp.

To explore how both reading and phonology skills could be related with these oscillation dynamics, Spearman correlations between the mean theta/delta ratio for each child for PC1 and both their phonological awareness and word reading ability were computed (Figure 1d). For dyslexic children only, a significant relationship was found for phonological awareness (*r*_s_ = −0.55, *p* = 0.027) but not for reading (*r*_s_ = 0.03, *p* = 0.916). This suggests a dyslexia-specific pattern of interplay between delta and theta power during natural speech processing that is progressively less atypical the better the child’s phonological awareness. To further control for PC1 sensitivity to developmental and reading level effects, typically-developing children were divided (as in [17]) into chronological age controls (same age as dyslexic children, better reading skill) and reading level controls (similar reading level but younger than dyslexic children) and tested as to whether they showed significantly different mean and variance on their theta/delta ratio. No significant differences were found across these subgroups (Figure S3). Correlations between PC1 theta/delta ratio and age; general IQ were also calculated. Once again, no significant relationships were found for any group (p > 0.05 for all correlations).

### Dyslexia-specific theta/delta power ratio differences for speech processing do not transfer to rhythmic syllable repetition but dyslexics show greater delta-band power

We then sought to understand whether these power modulation differences in dyslexia were specific to connected speech. To this end, we analysed a second EEG dataset recorded while different children with dyslexia and typically-developing children performed an audio-visual speech task without semantic or syntactic content (Figure 2a). No DLD children were recorded. In this paradigm, the child listened to trials comprising repetition of the syllable “ba” by a ‘talking head’ (see STAR Methods). Similar pre-processing pipelines and trial rejection rules as for story listening were applied. Age and neuropsychological profile of these groups is shown in Table 2 (see Table S2 for the full behavioural dataset).

**Figure 2.**
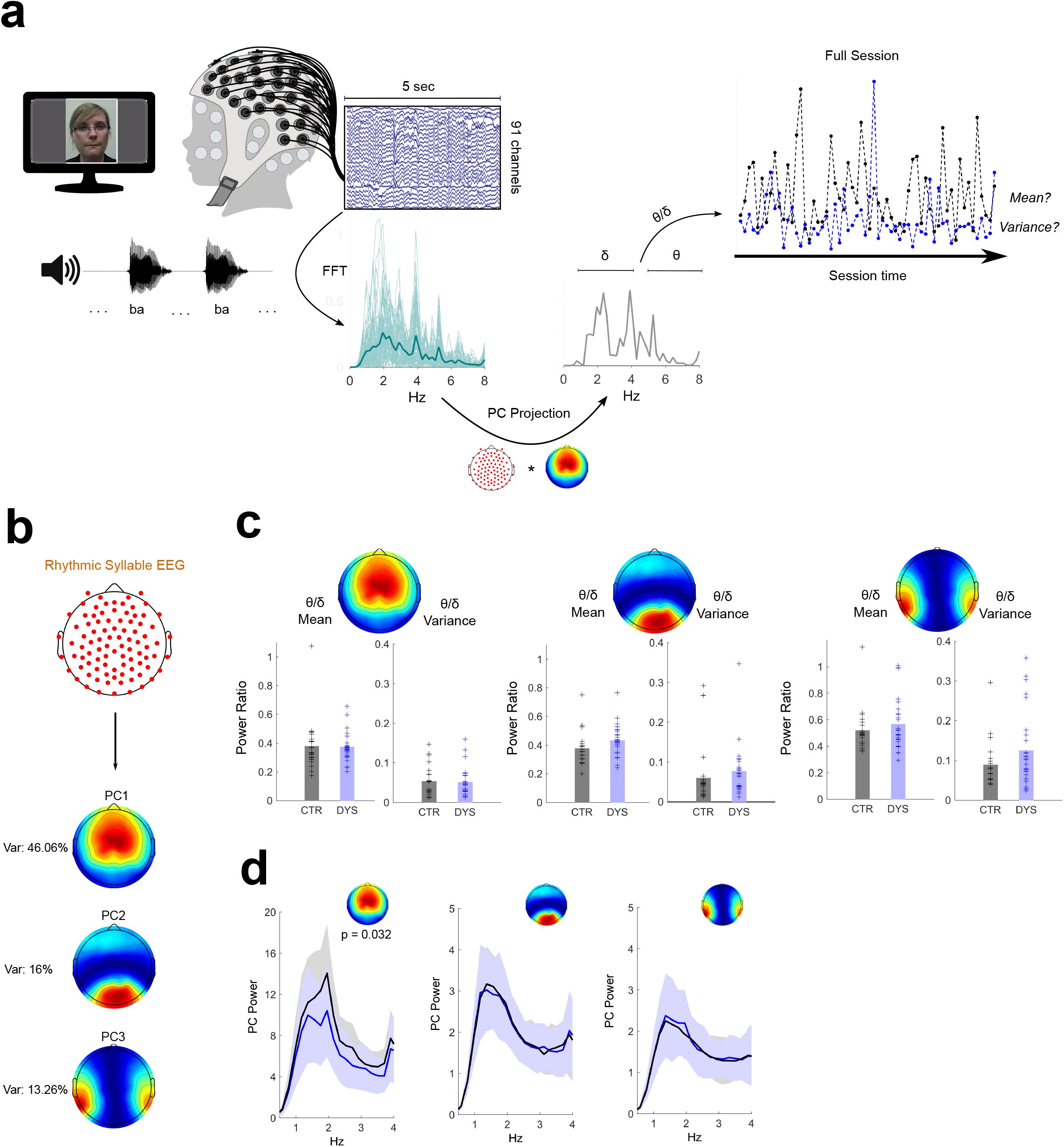
Dyslexic and typically-developing children exhibit similar theta/delta power ratio during rhythmic syllable listening. (a) Paradigm and pseudo-online analysis pipeline (see STAR Methods). EEG projection to PC1 was used for illustration. Sessions from a dyslexic (blue) and a typically-developing child (black) are shown. (b) Map of the three most important principal components – closely matching those of the story listening task (see Figure 1b). (c) No group differences were observed across all three PCs for theta/delta ratio mean or variance (CTR – typically-developing children, DYS – dyslexic children). (d) Typically-developing children show higher delta band power on PC1 only, which was maximal at the syllable presentation rate (2 Hz).

**Table 2.** Syllable repetition task: participant sample characterization

|  | **CTR** | **DYS** |
| --- | --- | --- |
|  | n = 21 (6F) | n = 27 (13F) |
| **Age (months)** | 109.1 (5.4) | 109.3 (6.8) |
| **PHAB_R<sup>a</sup>** | 102.6 (5.7) | 90.9 (11.5) |
| **TOWRE_W<sup>b</sup>** | 101.1 (7.7) | 79.4 (13.3) |
| **TOWRE_NW<sup>c</sup>** | 98.0 (8.6) | 76.9 (9.3) |
| **BPVS<sup>d</sup>** | 103.3 (11) | 103.7 (11.6) |
| **BAS_R<sup>e</sup>** | 99.5 (6.2) | 80.3 (7.4) |
| **BAS_S<sup>f</sup>** | 97.0 (6.1) | 78.6 (6.1) |
| **FSIQ<sup>g</sup>** | 104.1 (10.8) | 101.7 (9.9) |

CTR – Typically Developing (control group); DYS – Dyslexic children. Standard deviations in parenthesis
<sup>a</sup> Phonological Assessment Battery: Rhyme Awareness
<sup>b</sup> Test of Word Reading Efficiency: Words
<sup>c</sup> Test of Word Reading Efficiency: Non-Words
<sup>d</sup> British Picture Vocabulary Scale
<sup>e</sup> British Ability Scales: Reading
<sup>f</sup> British Ability Scales: Spelling
<sup>g</sup> Full Scale IQ (WISC-V subtests, see STAR Methods)
For all tests M = 100; SD = 15.

Once again, PCA was used to derive spatial filters for calculating theta/delta ratios in distinct brain regions. As with the story listening task, three PCs were sufficient to account for >70% of the variance (Figure 2b). Crucially, the spatial filters derived for both tasks shared remarkable similarities in terms of the spatial organisation and relative percentage of variance explained (compare Figure 2b to Figure 1b). The syllable repetition task showed a dominant central PC1 (total variance explained: 46%) followed by occipital (total variance explained: 16%) and bilateral temporal (total variance explained: 13%) PCs. The main difference between the syllable repetition and story listening task components was that the spatial configuration of PC2 in one task mirrored that of PC3 for the other task and vice-versa.

To investigate whether the rhythmic syllable repetition task would also show a group theta/delta ratio difference, group means and variances were compared. No group differences were found either for theta/delta mean or variance on any of the three dominant PCs (p > 0.05 for all comparisons) suggesting the theta/delta ratio effect is specific to connected speech (Figure 2c). In a further analysis for PC1, we found a significant group difference in delta band power (Z = 2.14, p = 0.032). Specifically, inspection of the EEG power spectra at 1-4Hz (Figure 2d, left panel) showed that the peak difference occured at 2Hz – the “ba” repetition rate – with a stronger response for typically-developing than dyslexic children. These data indicate a difference in neural steady-state responses to the rhythmic syllable targets. No other PCs showed this effect (Figure 2d, middle/right panels). Overall, these results suggest that differences regarding the interplay of delta and theta power dynamics in dyslexia do not transfer to a speech listening paradigm that lacks semantic / syntactic content or phrasal structure.

### DLD but not Dyslexic children show atypical delta-theta Phase-Amplitude Coupling during speech processing

Based on these results, we concluded that delta and theta power dynamics differed in dyslexic children (but not DLD) when compared to typically-developing children. We then questioned whether cross-frequency coupling of low-frequency oscillations – a neural mechanism crucial for sensory selection in speech processing [39] – would also differ across groups. Specifically, phase-amplitude coupling (PAC) differences involving the bands of interest were investigated using the story listening data. Principal Component Analysis was again used to spatially filter whole-brain data and delta-theta PAC was calculated for all children. Phase and amplitude of lower and higher frequencies, respectively, were computed using a time-frequency method that does not utilize bandpass filters, and a z-scored modulation index (zMI) was used to estimate neural PAC (see STAR Methods). For each participant, the average value and consistency (variance) of the coupling metric across the experimental session was calculated. The analysis pipeline is depicted in Figure 3a and mean / variance group comparisons of delta-theta PAC are shown in Figure 3b. While maximum coupling for most children occurred in the lower half of the delta band (1-2Hz) phase, the amplitude frequency for maximum PAC was highly variable across the theta band range (Figure S4). In Figure 3c (all panels) it is shown that this coupling occurs at a preferred delta phase of around ±π across the 3 dominant PCs for all groups.

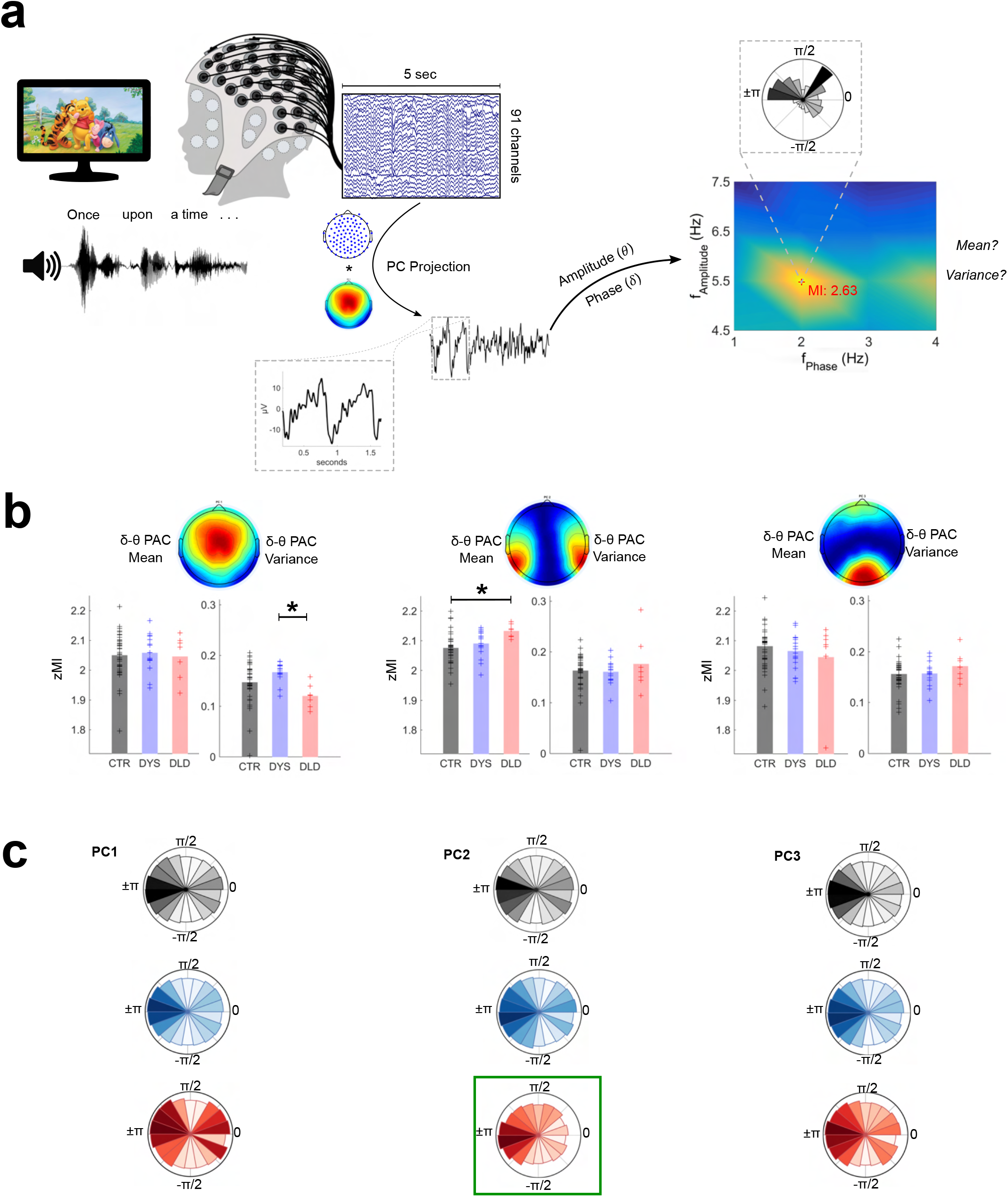

For PC1 (which had the largest channel weights in centrally-located electrodes), group differences were found for zMI variance (H = 12.1, p = 0.0024). Wilcoxon tests comparing typically-developing children and dyslexics (Z =-2.1, p = 0.036) or typically-developing children and DLDs (Z = 2.3, p = 0.022) were not significant following a Bonferroni correction. However, the two clinical groups showed significantly different zMI variances (Z = 3.3, p = 9.4e-4), with dyslexic children showing higher variance in maximum coupling compared to DLD across the experimental session (Figure 3b, left panel). Phase-amplitude plots (Figure 3c, left panel) indicate a relatively higher concentration of large amplitudes around the 0:π/4 phase of the delta band for DLD children. No significant differences were found in average zMI for PC1 across groups (p >0.05).

For PC2 (largest channel weights for electrodes covering temporal areas), differences were found across groups for mean zMI (H = 11.53, p = 0.0031), with DLD children showing higher delta-theta coupling (Z = −3.2, p = 0.0016) compared to typically-developing children (Figure 3b, middle panel). Phase-amplitude plots (Figure 3c, middle panel) showed that this significantly greater phase-amplitude coupling for DLD children stems from a higher concentration of larger theta amplitudes for the delta ±π phase bin, where larger amplitudes are also seen for the other groups. zMI mean comparisons between typically-developing children and dyslexics (Z = −1.5, p = 0.125) and between dyslexics and DLDs (Z = −2.2, p = 0.025) were not significant after a Bonferroni correction. Delta-theta variance of zMI for PC2 was similar across all groups (p > 0.05). No differences regarding maximum coupling strength across groups were found for PC3, the spatial filter with the largest weights on electrodes covering the occipital cortex (p > 0.05).

Finally, to test the sensitivity of PC1 and PC2 delta-theta PAC to developmental or reading level effects, once again we split the typically-developing children into subgroups of chronological age and reading level controls. No significant differences were found between these subgroups for delta-theta PAC (mean or variance) on either PC1 and PC2 (Figure S5).

Taken together, these results show that PAC metrics can distinguish children with DLD from both typically-developing and dyslexic groups. These differences in cross-frequency coupling were observed in principal components covering auditory / central areas. Similar to the dyslexia findings in theta/delta power ratios, no differences were observed in principal components covering primary visual areas.

### Supervised temporal-spatial EEG filters reveal oscillatory power differences across experimental groups

The theta-delta ratio and PAC results suggest that dyslexic and DLD children show distinct atypical patterns of low-frequency oscillatory dynamics during natural speech listening. However, potential group differences in spatial patterns regarding each individual oscillatory frequency have not yet been addressed. We therefore returned to the story listening data and sought biomarkers that could show group differences across delta, theta and beta oscillations. Previous Brain-Computer Interface studies have used Common Spatial Patterns (CSP, see STAR Methods) to find optimal EEG patterns successfully as well as decode brain states [40] and silent speech [41]. Applied to the present study, this linear method finds supervised filters (using labelled data) that simultaneously maximize the EEG signal variance for one group of children while minimizing the variance for the other, and vice-versa (Figure 4a). To uncover spatial filters that would discriminate between our 3 groups, 3 sets of CSP filters were calculated for delta, theta and beta oscillations respectively (translating to a total of 9 sets). Specifically, we compared 1) typically-developing versus dyslexic children, 2) typically-developing versus DLD children, and 3) dyslexic versus DLD children for each oscillatory band. Each CSP calculation yielded the same number of filters as the number of recorded channels. As in previous BCI research (see [40] for an overview), only a subset of these filters was analysed. For each group comparison, the 2 spatial filters maximizing the variance for each group (i.e., for *m* = 2; total number of filters = 4) were computed and participant differences were assessed. Twelve comparisons were made (4 spatial filters x 3 brain rhythms) for each group combination and only the filters showing significant differences in non-parametric ANOVAs that survived Bonferroni correction were considered (threshold p-value = 0.05/12 = 0.004).

**Figure 4.**
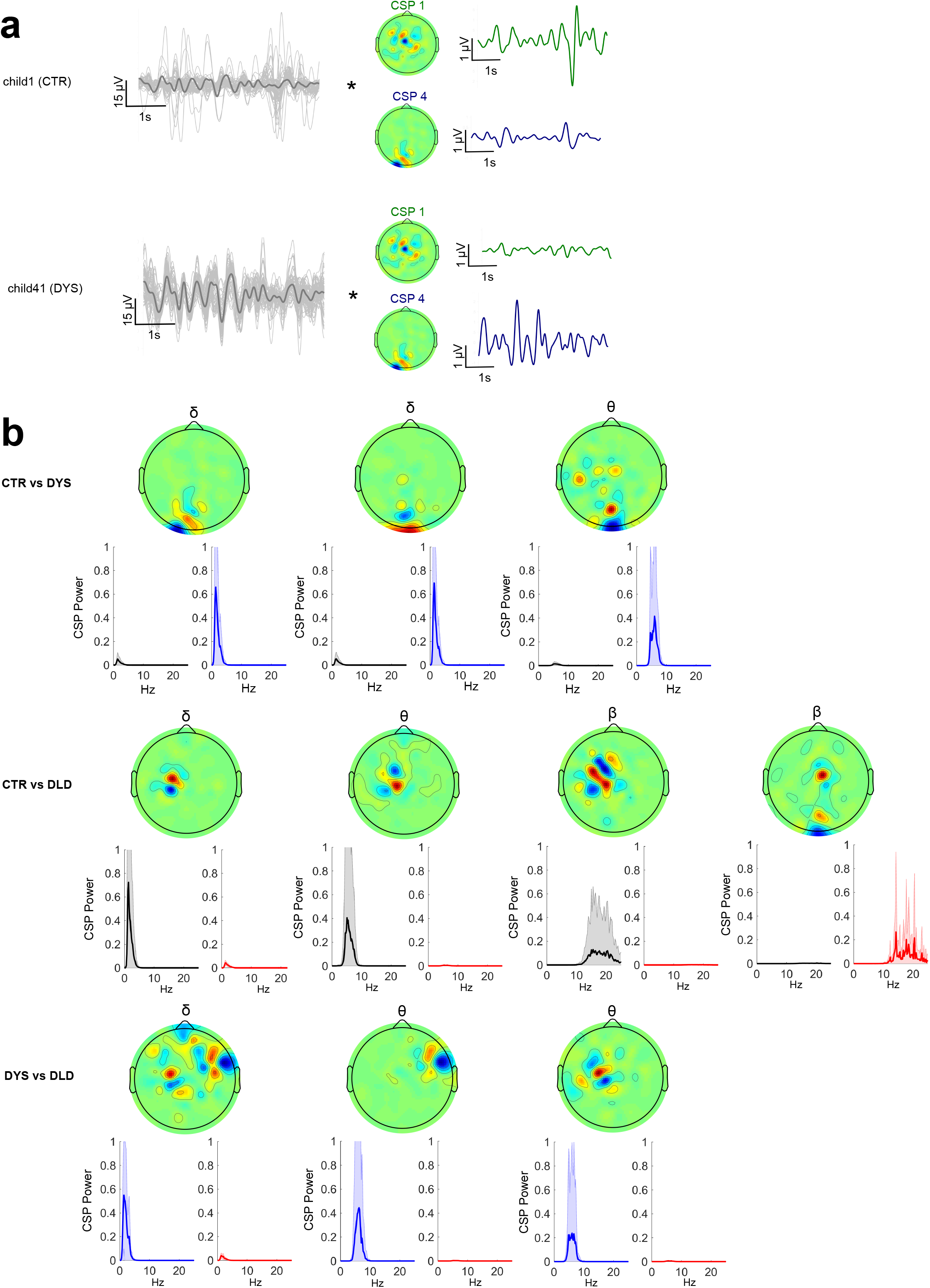
Common Spatial Patterns (CSP) enable discrimination between dyslexic, DLD and typically-developing children during speech listening. (a) Spatial filtering applied to the original EEG epochs (each channel depicted in grey, average in black). Spatial filters allow discrimination of EEG signals from experimental groups in the story listening task by selectively maximizing the signal variance for one group while minimizing for the other. (b) CSP filters for delta, theta and beta showing average power for each group for each filter. These CSPs showed significant differences at the group level (p < 0.05, Bonferroni corrected). Upper panel shows filters yielding significant group differences for the typically-developing children vs dyslexic, middle panel for the typically-developing vs DLD children and lower panel for the dyslexic vs DLD children.

Figure 4b shows the results. When comparing typically-developing children with dyslexic children (Figure 4b, upper row), filters relying mostly on occipital channels were the most discriminative and maximized the EEG power for dyslexic children on delta and theta rhythms. When comparing typically-developing children with DLD children (Figure 4b, middle row), central / left lateralized CSP filters maximized the EEG power for typically-developing children (minimizing for DLDs) across all 3 brain rhythms. A spatial filter focusing on occipital channels minimized the variance for typically-developing children while maximizing variance for DLD children in the *beta band* range. Overall, strongly occipital EEG filters minimized the EEG power of typically-developing children while maximizing power for both DLD and dyslexic groups (see top and middle rows, Figure 4b). Finally, when comparing dyslexic with DLD children (Figure 4b, bottom row), the significant spatial filters were on delta and theta rhythms. These CSPs maximized the EEG power for dyslexics and had strong weights on the right temporal and left central channels.

### Supervised temporal-spatial filtering using CSP enables epoch-based classification of dyslexia on a story listening task

Despite the significance of between-group differences for CSP-based biomarkers, it is not yet clear whether features based on these CSPs have enough robustness to be used in a classifier – especially for the 5-second window epochs employed here. To test this, we cross-validated linear Support Vector Machines – with delta, theta and beta filtered EEG CSP feature inputs, respectively – using Leave-One-Subject-Out crossvalidation. This method allows repeated assessment of the model’s performance without leaking subject-specific information to the training set (see Figure 5a and STAR Methods). SVMs have been frequently used as classifiers in the EEG literature (see Ref. [38] for a recent example) and have shown performances ranging from 0.6-0.95 AUC in EEG classification problems using longer inputs. Each classifier was trained with a different number of CSP filters to detect the optimal number of features. Importantly, for each cross-validation fold, CSPs were calculated only on the training set to avoid feature information leakage from the test set and, consequently, overfitting. To avoid “double dipping”, no information from previous group analyses was used to influence the choice of specific spatial filters (see STAR Methods).

**Figure 5.**
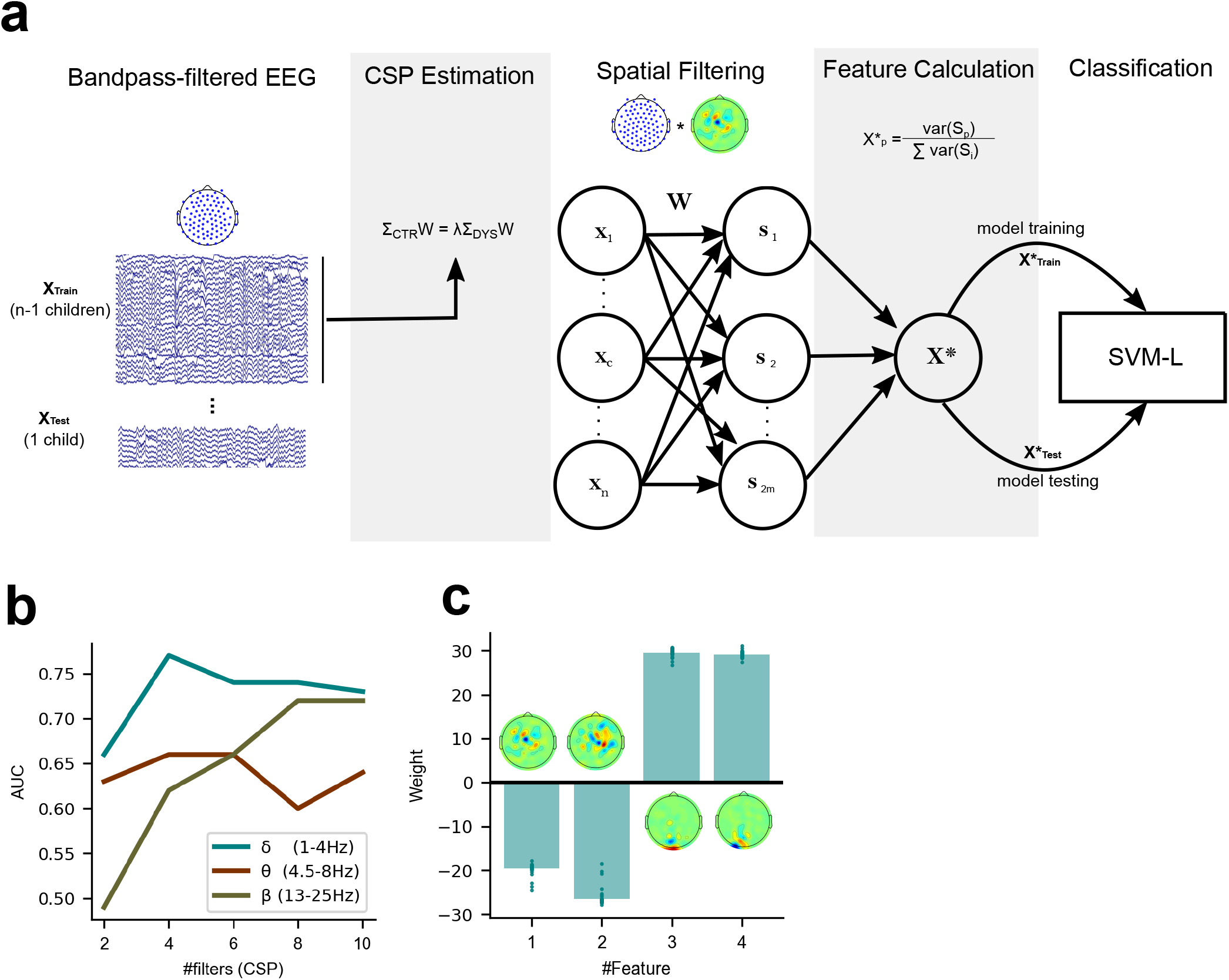
Classification of dyslexia using a linear classifier trained with features based on Common Spatial Patterns. (a) Leave-one-subject-out cross-validation pipeline for the Linear SVM algorithm (total number of features = 2m). (b) Performance (based on the area under the curve – AUC) difference between linear classifiers using delta, theta and beta-CSP filters across a different number of features (#filters). (c) Delta-CSP feature weights of the linear classifier are stable across cross-validation folds.

The receiver operating characteristic (ROC) curves for the classifiers are presented in Figure 5b. Simple linear SVMs using 4 feature variables (i.e., spatial filters) with the delta-CSP classifier reached an area under the curve (AUC) of 0.77. Since this delta-CSP classifier showed the best performance (compared to all the theta- and beta-CSP classifiers, see Figure 5b), we further analysed the spatial location of its most important CSP weights, the relationship between these CSPs and EEG activity, and the variance of the classifier’s weights for the CSP features across all 56 folds (defined as the number of participants in Leave-One-Subject-Out cross-validation).

The results showed that the two negative-weighted CSP features were those that maximized the signal variance for typically-developing children and minimized the variance for dyslexic children (Figure 5c). These spatial filters were more strongly weighted on central / right lateralized channels and showed relatively higher correlations with channel activity in left/frontal and right/central areas of the scalp for control children when compared to dyslexic children (Figure S6a,b). On the other hand, the largest positive weights were the two spatial filters that maximized the signal variance for dyslexic children and minimized the variance for typically-developing children. These filters were mainly focused on occipital channels, and the absolute values of their weights were similar to those of other spatial filters. These occipital CSP filters showed stronger correlations with electric potentials in larger areas of the scalp for dyslexic children (versus typically-developing children), extending to parietal and central electrodes (Figure S6c,d). Feature weights remained stable across the crossvalidation process (Figure 5c), despite CSP filters being slightly different for every fold (given the changes in the training set). CSP channel weight estimations across folds were also highly consistent (Figure S7).

Taken together, these results show that dyslexia classifiers can be based on EEG-CSP features. Furthermore, in line with our prior studies [14, 15, 42, 43], delta-band features were found to be most useful to identify developmental dyslexia. In contrast with our earlier findings regarding oscillatory dynamics (theta/delta ratio), we find that occipital channels play an important role regarding group discrimination. The importance of EEG delta rhythms in story listening tasks for children with dyslexia converge with recent MEG speech-brain coherence studies [44, 18].

### Rhythmic syllable repetition CSP provide useful features for dyslexia classification on the story listening task and vice-versa

Given the apparently central role of delta-band responding for dyslexia, we then investigated whether these same 4 delta-band spatial filters that identified dyslexia in story listening EEG might contain useful information for classifying dyslexia in the rhythmic syllable repetition EEG. We thus trained a linear SVM classifier to predict dyslexia on the rhythmic syllable repetition dataset, using 4 delta-CSP filters trained using the story listening dataset (Figure 6a). To obtain baseline performance for a classifier trained on the most discriminative CSP filters computed for the syllable repetition task, a pipeline similar to the story listening task (see Figure 5a) was applied to this dataset.

**Figure 6.**
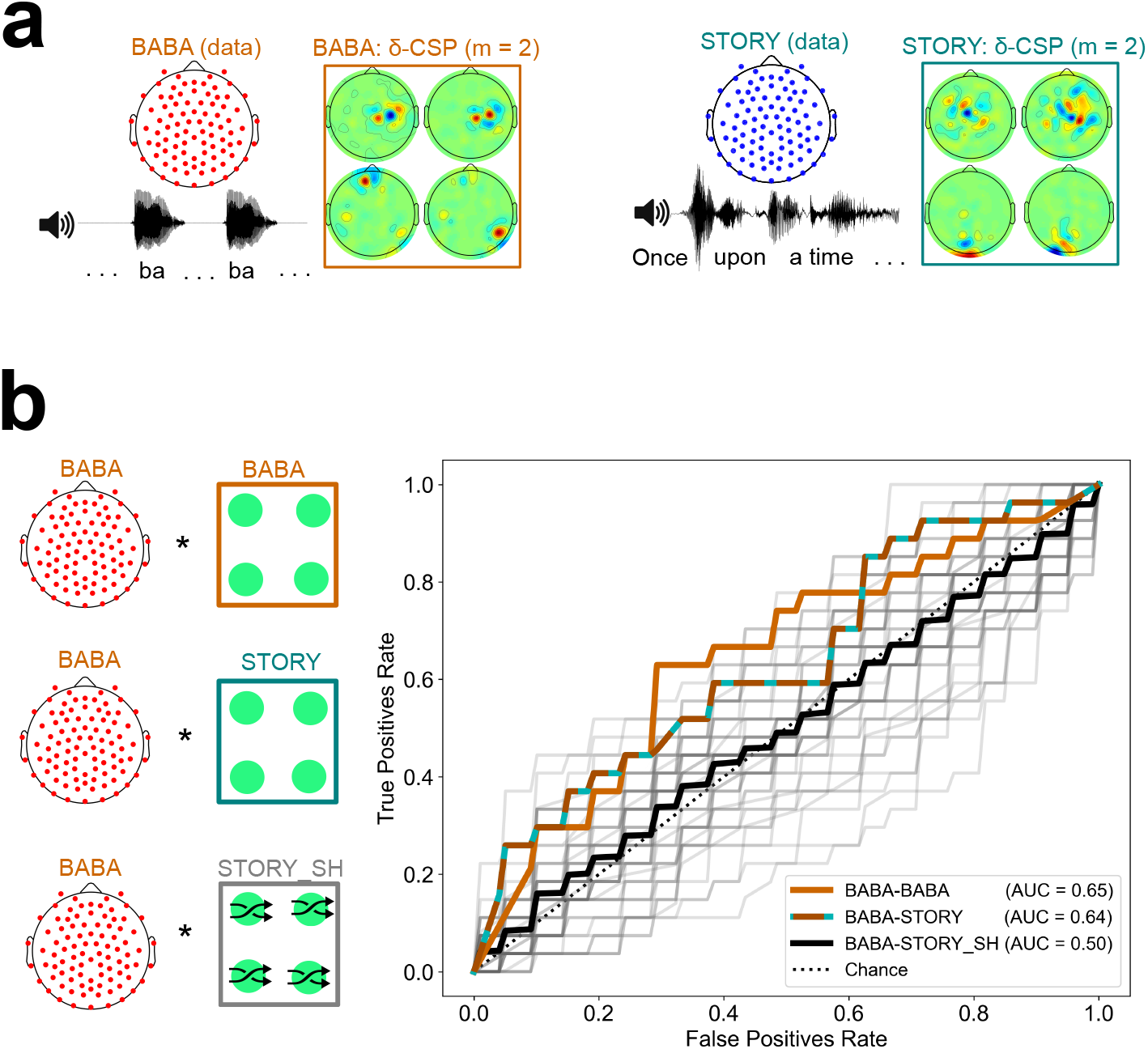
Analyses revealing successful transfer-learning of delta-CSP features for dyslexia classification across story listening and rhythmic syllable repetition tasks. (a) Delta-CSP filters for the rhythmic syllable task (orange) show higher weights on right temporal channels when compared to the delta-CSP filters of the story-listening task (teal) that show higher weights on occipital / central channels. (b) Similar AUCs for linear SVMs classifying dyslexics and typically-developing children on the rhythmic syllable repetition task with its original cross-validated CSPs (BABA-BABA) and with CSPs derived from the story listening task (BABA-STORY). SVMs using spatially shuffled versions of the story CSPs (BABA-STORY_SH) resulted in chance-level performance (black).

Figure 6b shows the results. SVM performance regarding classification of dyslexic children on the rhythmic syllable repetition data with the original CSP filters versus the story-listening CSPs was very similar (0.64 and 0.65 AUC, respectively). However, this generalization may occur simply because temporal filtering (i.e., the 1-4Hz bandpass for delta) is the sole provider of a whole-brain metric for classification, irrespective of spatial filter weights on specific channels. To test this possibility, we trained 50 linear SVMs, each one using a different random permutation shuffle of the story-listening CSP weights (BABA-STORY_SH condition, see also Figure S8). The average AUC performance of these classifiers was at chance level (black ROC curve, AUC = 0.5), and the model using the original story-listening CSPs was better than 96% of models using its shuffled versions at predicting dyslexia status from the rhythmic syllable repetition task (Figure 6b, grey ROC curves). This result showed that the specific configuration of story-listening EEG spatial filters is encoding important information for classifying dyslexia using the rhythmic syllable repetition EEG. Accordingly, speech rhythm processing may be the common factor being identified by these classifiers regarding dyslexia and potential biomarkers.

## Discussion

Here we show that different automatic low-frequency neural oscillatory responses to connected speech in a passive story listening task can uniquely identify children with two developmental disorders of language with distinct cognitive profiles: DLD (syntax impaired) and dyslexia (phonology impaired). Further, we have shown that low-frequency oscillatory activity during speech listening can reliably classify children with dyslexia. These data suggest that mechanistic relationships between low frequency (i.e., delta and theta) oscillatory bands are fundamental to understanding the aetiology of dyslexia and DLD, as predicted by TS theory [10, 45]. The data also show the importance of studying children when trying to understand causal factors in developmental disorders of learning. Adult studies have assumed that faster-rate (phonemic, >30Hz) speech envelope modulations should be impaired in individuals with dyslexia in alphabetic orthographies, as grapheme-phoneme conversion is fundamental to reading proficiency [46]. However, developmental studies show that phonemic information is learned via *reading experience,* with fast-rate oscillations showing atypical patterns in children only *after* the onset of reading [47,44], and minimal gamma-band synchronisation present at the beginning of the dyslexic reading trajectory [44]. Further, it has been shown that low-frequency oscillations are involved in phoneme-level processing of speech [48, 49], and that atypical speech entrainment in the right hemisphere for low frequencies can be associated with phoneme-level processing in dyslexia [17]. We expect that the atypical oscillatory responses found here for dyslexia and DLD may generalize across languages.

Low-frequency oscillations usually signal communication involving large populations of neurons in relatively large brain areas, while higher frequency oscillations are nested in these rhythms and act more locally [50]. The potential EEG markers found here for dyslexia and DLD were indeed spread across large areas of the scalp. The direct consequence of such nesting is that atypical oscillatory patterns at slower frequencies (observed here in both dyslexia and DLD) may have downstream effects regarding the magnitude of oscillatory responses in high frequency brain rhythms. Therefore, our data are compatible with previous adult dyslexia studies indexing atypical fast-rate oscillatory power [46] for non-speech steady-state stimuli. The loci found here for atypical low-frequency oscillatory responses are also in line with clinical observations of dispersed structural brain abnormalities in dyslexic participants [51] as well as molecular differences in genes that regulate the development of the entire language network in both DLD and dyslexia [52–54].

A key finding in our study is that the *relative* oscillatory power of theta/delta responses is atypical in dyslexic children during a story listening task. Such differences were seen for an electrode ensemble strongly weighted around the centre of the scalp. These loci are consistent with the hypothesis that children with dyslexia are accessing a mental lexicon that contains atypical auditory representations with less accurate representation of delta-band speech envelope information [15]. Cognitive processing of prosodic information is impaired in children with dyslexia when compared to both age-matched and reading-level matched (hence younger) controls [13, 55]. In line with these prior behavioural findings, we find this oscillatory marker to be associated with performance in phonological rather than reading measures. These atypical patterns of neural processing are likely to influence the mental representation of prosody in language from infancy and throughout development [56].

We found that children with dyslexia no longer showed a difference regarding the theta/delta oscillation power ratio compared to typically-developing children for a rhythmic syllable repetition task. In prior work, this rhythmic task revealed a different preferred phase in the delta band in the dyslexic child brain [14, 15, 43]. Here, children with dyslexia showed significantly lower oscillatory delta band power in this task, with maximal differences at the syllable presentation rate (2Hz). Given the audio-visual nature of the task, this finding suggests that steady-state responses from the dyslexic brain are weaker even when the task allows supra-additive responses due to congruent information across visual and auditory modalities [34, 57, 58]. This is in line with prior research showing poorer conversion of lip-read information into auditory speech representations for dyslexic participants [59].

We further observed atypical neural cross-frequency phase-amplitude coupling for delta-theta in DLD but not in dyslexia (story listening task). This effect was observable on electrode ensembles covering bilateral temporal regions, suggesting that sensory representation / binding of sound features is affected in children with DLD. This is in line with studies showing atypical auditory sensory processing in DLD regarding many non-speech acoustic parameters, including amplitude rise times. One prior study found that children with both disorders were significantly poorer than typically-developing children in recognising sentences presented as 4-channel or 8-channel vocoded speech, a method of degrading speech that increases reliance on the speech envelope [60]. An unusually strong neural dependence of delta-theta coupling to process speech could potentially explain these difficulties for the DLD children.

Our CSP filtering approach revealed that filters with larger weights on channels covering occipital areas consistently showed group differences between typically-developing children and both dyslexic (delta and theta bands) and DLD children (beta band), with the clinical groups showing notably high oscillatory activity in occipital regions. While occipital areas are not typically associated with acoustic/phonological deficits, these findings are compatible with a previous developmental literature suggesting compensatory visual mechanisms accompanying auditory processing deficits in children with dyslexia [61], as well as atypically strong sustained delta oscillations in dyslexic children in posterior brain regions while processing phonological and semantic information [62]. These differences may also pertain to an oscillatory footprint for the visual word-form area (VWFA) that is activated during receptive language processing by adults [63] – along with previously reported functional changes regarding the VWFA for children with dyslexia [64, 65]. DLD children showed weaker oscillatory activity across both delta and theta rhythms when CSPs with strong weights for left-lateralized temporal and central channels were used. In adults, delta and theta rhythms have been linked to processing acoustic “edges” (amplitude rise times) in speech [20, 66]. Meanwhile, the magnitude of beta oscillatory responses has been associated not only with comprehension but also with predictive coding of speech [67]. Taken together, these findings may suggest that speech processing and speech prediction may be atypical in DLD children, with a clearly different phenomenological manifestation from that observed in dyslexia.

Large non-linear EEG-based classifiers have previously been engineered for dyslexia using long time windows of AM noise, with degrees of success reaching an AUC ~ 0.8 [37, 38]. By contrast, we test classification performances for both dyslexic and typically-developing children using a linear classifier and short epochs of naturalistic speech listening data. We find features from a minimal number of EEG spatial patterns for delta oscillatory responses, which show high-magnitude differences between dyslexic and neurotypical children. We then trained a classifier for the story task using CSPs from the less naturalistic rhythmic syllable repetition task enabling transfer-learning across datasets. Crucially, the training of these classifiers showed comparable performances to their original CSP features. This suggests that we are picking up oscillatory patterns related to general speech rhythm processing. Interestingly, half of the CSPs from the syllable repetition task were specifically located in right-lateralized temporal areas. This indicates differences in the symmetry of spatially filtered delta oscillations between dyslexic and typically-developing children, matching adult data [68, 69].

In conclusion, we have identified relationships between low-frequency EEG oscillations related to different neural speech processing mechanisms that are selectively atypical in dyslexia versus DLD. Further, we find that the magnitude of delta oscillations in a story listening task shows a consistently different pattern between dyslexic and typically-developing children, potentially enabling the development of a generalizable classifier for developmental dyslexia. Our cross-dataset approach provides evidence that these oscillations are likely related to speech rhythm processing, a core tenet of TS theory. We also demonstrate transfer-learning of EEG features for identification of children with dyslexia across different receptive speech tasks and different samples of children. Taken together, our data provide robust evidence for a temporal sampling deficit in two developmental disorders of language learning, dyslexia and DLD [10, 45].

## STAR Methods

### Participants and Experimental Paradigms

Two EEG datasets generated in prior studies by our group were used for the modelling [17, 43]. The [17] data set was collected during a story-listening task, while the [43] data set was collected during a rhythmic syllable repetition task (listening to the syllable “ba” repeated every 500 msec). In each case, cortical activity was recorded using the scalp potentials measured by non-invasive EEG. In the story listening paradigm, participants were presented with an audio-story for 9 minutes read by a female Australian English speaker while EEG was recorded (Fig. 1a). The stimulus was presented monophonically at a sampling rate of 44100⍰Hz using loudspeakers in a silent room. Participants also watched a cartoon corresponding to the story (Winnie the Pooh), but the visual input was not synchronized to the detailed temporal events coming from the auditory modality (i.e., to the speech). EEG was recorded using 129-channel Hydrocel Geodesic Sensor Net (HCGSN), NetAmps 300 amplifier and NetStation 4.5.7 software *(EGI* Inc.). The sampling rate of this system was 1⍰kHz and channel impedances were always kept below 50 kΩ throughout the session.

The second paradigm was a rhythmic entrainment task [70]. Children listened to rhythmic speech comprising multiple repetitions of the syllable “ba” at a 2Hz rate. In common with the story-listening task, the auditory targets were presented at 44100 Hz, albeit via in-earphones. Synchronized visual information (a “talking head” providing articulatory cues onsetting 68ms before the syllable onset) was also presented (synchronized audio-visual task). Participants were instructed to concentrate their gaze on the lips of the talking head to prepare for every new trial. In each trial, the syllable “ba” was presented 14 times and the child was instructed to press a key on the keyboard if they detected any syllable that violated the uniform 2Hz rhythm. Feedback was presented after each 14-item trial. The full experimental session consisted of 90 trials divided into 3 blocks of 30. Each block comprised 25 trials where a rhythmic violation would occur randomly between the 9^th^ and 11^th^ syllable and 5 catch (i.e., absence of rhythmic violation) trials. For the rhythmic violation trials, the degree to which the violator was out of sync changed with each child’s performance on the task to optimize their level of engagement. This was achieved by a three-down one-up staircase procedure – if a child correctly identified 3 violations in a row, the stimulus onset asynchrony would be reduced 16.67ms on the following non-catch trial and if a violator was not detected, the deviation would be increased 16.67ms. Similar to the story-listening set-up, a 129-channel Hydrocel Geodesic Sensor Net was used to record EEG scalp potentials during the task. The sampling rate was 500 Hz and electrode impedances were kept below 50 kΩ.

For the current paper, data from 65 Australian English children who received the story listening paradigm and data from 48 British English children who received the syllable repetition task were used (see Supplementary Tables 1 and 2 for tabulation of the children). The status as typically-developing/dyslexic/DLD for the Australian children was ascertained using a range of neuropsychological and cognitive tests. The children included had non-verbal IQ scores within the normal range (85 and above) on the Kaufman Brief Intelligence Test (KBIT, [71]). To qualify as dyslexic, children had to score at least 1 SD below the norm of 100 for single word or nonword reading measures on the TOWRE (84 and below, Test of Word Reading Efficiency, [72]), and show average scores (less than 1 SD from the norm of 100, so 85 or above) on at least one of the measures of language development, the TROG (Test of Receptive Oral Grammar, [73]), CELF (Clinical Evaluation of Language Fundamentals, [74]) or WIAT Vocabulary [75]. Children also received the CTOPP (Comprehensive Test of Phonological Processing [76]) as a measure of phonological awareness, a test requiring the oral blending or elision of words, syllables and other phonological units. To qualify as DLD, children had to show standard scores at least 1 SD below the norm of 100 for the TROG, WIAT and CELF measures, and an average score (less than 1 SD from the norm of 100) on the reading measures. Please note that the vocabulary test was changed during the project from the CELF vocabulary scale to the WIAT vocabulary scale, see note in Table S1. Some children were borderline regarding the diagnostic criteria but were still included following discussion by the research team (8 participants, CTR 23, 26, 29, DYS 1,14, and DLD 4,6,7 in Table S1). Fewer DLD children (N=7) were in the Australian sample than dyslexic children (N=16). No participants with ADHD were included. After the pre-processing pipeline, 2 typically-developing participants were excluded due to noisy EEG measurements so that, in the end, data from a total of 63 Australian participants were analysed.

For the British children, typically-developing versus dyslexic status was also ascertained by neuropsychological and cognitive testing (see Supplementary Table 2). Participants had full scale IQs (FSIQ) in the normal range (85 and above) as estimated from four subtests of the Wechsler Intelligence Scale for Children (WISC [77], similarities, vocabulary, block design and matrix reasoning [78]. To qualify as dyslexic, children had to score at least 1 SD (15 standard points) below the standard score of 100 on at least two of 4 measures of single word or nonword reading and spelling (scoring 84 or less on the British Ability Scales, BAS [79], and TOWRE [72]). Children also received the Phonological Awareness Battery (PhAB, [80]) rhyming test as a measure of phonological awareness. One child was borderline regarding the diagnostic criteria but was still included given their poor phonology (DYS 9). Where test scores were missing for reasons such as school absence, standard scores from the next nearest test session are shown (affects DYS 7, 9, 22). 21 British typically-developing children and 27 dyslexic children participated. As the aim of the current modelling was to distinguish neural characteristics of the dyslexic versus DLD brain, the typically-developing children covered a range of ages, language and reading levels.

### EEG signal pre-processing

The pre-processing pipeline was similar for both the story listening and rhythmic syllable repetition data to ensure analysis consistency and maximize the validity of cross-set comparisons. The EEG signal was first bandpass-filtered between 1-25Hz using a 4^th^ order Butterworth filter to remove very low-frequency and high-frequency noise (including power line noise). A zero-phase filtering method (backwards and forwards filtering – using the *filtfilt* function in MATLAB) was used to prevent phase shifting and reconstruct the original properties of the signal of interest as much as possible. Data were subsequently re-referenced to mean mastoids. Since one of the objectives of this study was to classify signals in a window suited for Brain-Computer Interface (BCI) applications – whether for potential diagnosis or operant learning interventions – data was epoched to 5-second windows. The epoching of the story-listening data in short windows not only provides a way to evaluate the behaviour of any EEG signal metric across the session but also allows for the comparison with the rhythmic syllable trials that were epoched in time windows of similar length (5 seconds). To further remove noise sources like blinks or EMG activity, for each trial, channels with voltages over the absolute value of 100 μV were considered noisy and were interpolated using the spline method (EEGLAB; [81]). If over a third of the total number of channels were considered noisy, the trial was excluded. As in [17], data were downsampled to 100Hz to reduce processing time and memory requirements, and EEG electrodes positioned at the jaw, mastoids, and forehead were removed from the analysis (91 channels total; see Figure 1b). After the pre-processing pipeline, each participant kept on average 88.63 epochs (st.d = 29.3) for the story listening task, and 45.23 epochs (st.d = 18.61) for the rhythmic syllable task.

### Deriving unsupervised spatial filters using Principal Components Analysis

Principal Components Analysis (PCA) is used in this work to 1) allow dimensionality reduction of a 91-dimensional space of channels and in the process 2) create relevant spatial ensembles of channels that represent distinct (uncorrelated) sources of cortical activity. This ensemble analysis aims to create a more meaningful basis for analysing the whole-brain EEG signal than just relying on multiple single-channel analyses. In general, PCA will find the set of vectors *A* that project the original data *X* into a new space explaining the maximum original uncorrelated variance in the least number of vectors possible. This is one of the bottom-up methods for deriving spatial filters for EEG in an unsupervised manner (i.e., without looking at group labelled data). PCA can be conceptualised as an eigenvalue problem:

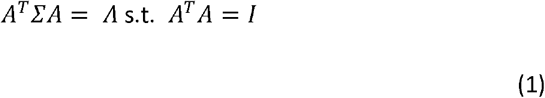

where Σ = *X^T^X*. In practice, the matrix *A* was estimated using Singular Value Decomposition *(svd* function in MATLAB) and consists of a set of eigenvectors sorted by their eigenvalues (diagonal elements of matrix *Λ*). The data *X* consisted of the concatenation of standardized *t* epoch segments 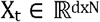 (where *d* is the number of channels and N is the number of datapoints on an epoch) along the second dimension so that 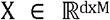 with *M = total number of trials * N*. The relative channel weights for each PC were determined simply as the square of their original coefficient for each *a* ∈ *A*, since ∑*a*^2^ = 1. Initially, to decide on how many principal components to retain (i.e., the total new dimensionality of the data), a threshold of 70% variance explained was used. 3 PCs were found to be sufficient in both the story listening and the rhythmic syllable repetition EEG data (Fig. 1b). Additionally, a further analysis of the scree plots (Figures S1a, S2a) revealed minimal increases of variance explained after PC3 which would usually explain 10 times as much variance as the next best PC on both datasets. A quick look at the channel weights of the first 10 PCs (Figures S1b, S2b) suggests the first 3 PCs for the story listening and rhythmic syllable repetition EEG datasets are less scattered and more biologically plausible when compared to the other 7 PCs. Taken together, this evidence provided additional confidence in choosing 3 as the optimal number of PCs to retain.

### Principal Component band power ratio

To calculate the band power ratio for every PC band, Welch’s method was used first to estimate the broadband power spectral density of each epoch projected in each PC. A single Hanning window covering the epoch’s full length (500 datapoints at a sampling frequency of 100Hz) was used. In practice, this was calculated using MATLAB’s Welch method implementation in the *pwelch* function. Each band power was calculated by averaging the discrete Fourier transform points belonging to each frequency band interval (delta: 1-4Hz theta: 4.5-8Hz). The theta/delta band ratio was calculated by dividing the averaged band power of theta and delta for each epoch. The band ratio metrics for each subject were calculated by taking the low order statistics (mean and variance) of their epochs’ theta/delta ratios across the session.

### Phase-Amplitude Coupling (PAC)

To estimate phase and amplitude for each epoch on the PAC analysis we relied on a recently published time-frequency approach that does not rely on bandpass filtering [82–84]. Indeed, bandpass filtering artefacts, such as approximating the filtered signal to a sinusoid, especially in small bandwidth and high order filters, can be problematic for PAC estimation. Note only does the time-frequency approach used here solve this problem, but it also has interesting properties such as high frequency resolution and is more robust to noise, different data lengths and sampling rates [82]. The method relies on the complex time-frequency distribution based on the interaction of energy at frequency *f* at a given time *t* known as Reduced Interference Distribution (RID) – Rihaczek distribution. The RID-Rihaczek distribution is a modified version of the Rihaczek distribution that uses the Choi-Williams kernel to filter out the cross-terms:

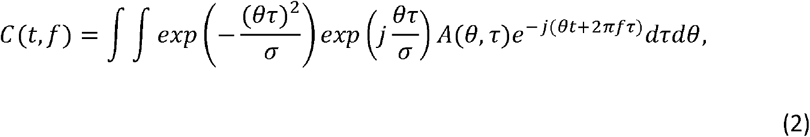

where 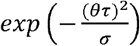 is the Choi-Williams kernel, 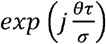 is the kernel function of the Rihaczek distribution and *A*(*θ*, *τ*) is the ambiguity function of the given signal *x*(*t*). This distribution belongs to Cohen’s class of distributions, reflecting the time-varying energy and phase of the signal. For an analytic signal *x*(*t*) = *A*(*t*)*e^jϕ(t)^* with Fourier transform *X(f)* = *B*(*f*)*e^jθ(f)^* the instantaneous amplitude (for higher frequency) and phase (for lower frequency) based on the Rihaczek distribution are given by:

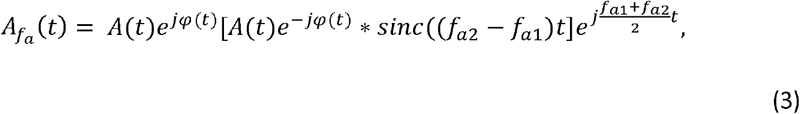

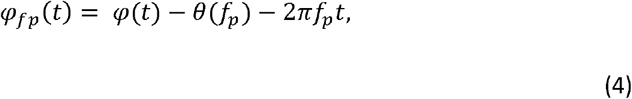

where f_a1_ and f_a2_ define the high frequency amplitude bandwidth and ⍰(t) and θ(fp) refer to the phase of the low frequency band in the time and the frequency domains, respectively. In practice, the calculation of the amplitude and phase for each epoch was performed by using the MATLAB code provided in the original publication [82].

The phase-amplitude coupling metric is then calculated using the Modulation Index (MI) [85]. This method seems to be relatively robust to phase biases [86] compared to other methods such as the mean vector length [87] and quite conservative in low signal-to-noise ratio conditions. MI discretizes the phase angle time series of the phase frequency into *N* phase bins and computes the average power of the modulated frequency for power in each bin *j*. In this work, we set *N* = 18, similar to that used in other studies (i.e., 20º wide bins). Coupling is operationalized in an information-theoretical way as the deviation of the phase-amplitude histogram from the uniform distribution:

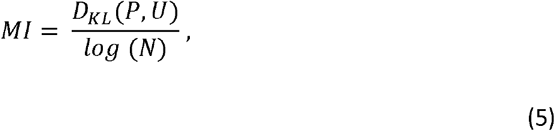

where *N* is the number of bins, D_KL_ is the Kullback-Leibler distance between the phase distribution *P* and the uniform distribution *U*:

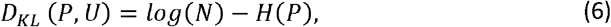

and *H* is the Shannon Entropy of the phase distribution:

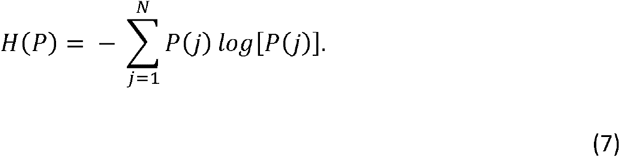

For each epoch, MIs were calculated across the full bandwidth of phase and amplitude bands. For each epoch, MI values across the comodulogram were z-scored (zMI) and the maximum zMI was the PAC metric for every epoch. Similar to the band ratio analysis, the average and variance of epoch PACs was taken for every subject. In practice, zMI was applied using a custom-made Python code (which will be made available upon request).

### Deriving supervised spatial filters using Common Spatial Patterns

The Common Spatial Patterns (CSP) algorithm [40, 88–90] has some similarities with PCA in the sense that it is also an eigendecomposition method. However, instead of finding the filters that maximize uncorrelated signal variance like PCA, its aim is to find filters that maximize the variance for one group of subjects and minimize the variance for the other (i.e., discriminative EEG spatial patterns). Therefore, CSP is a supervised method that calculates spatial filters based on labelled data. Given a set of *t* epoch segments 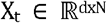 (where *d* is the number of channels and *N* is the number of datapoints on each epoch), epoch covariances 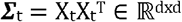, and *Σ_1_* and *Σ_2_* as the average epoch covariances for group 1 and group 2 subjects, CSP is calculated by the simultaneous diagonalization of the two average covariance matrices

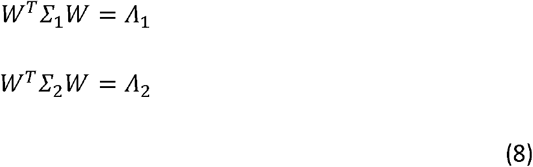

where W is commonly determined so that Λ_1_ + Λ_2_ = I (with Λ being a diagonal matrix of eigenvalues). Technically, this is achieved by solving the generalized eigenvalue problem

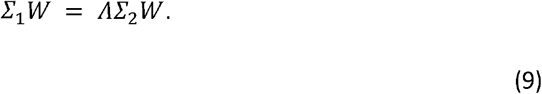

In practice, this was calculated either using MATLAB’s *eig* function or Python *linalg.eigh* function from the *scipy* package. The spatially filtered signal S of this set of EEG epoch segments is then given by

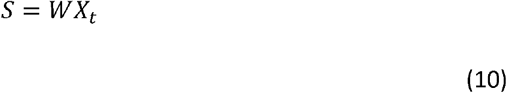

with the leftmost spatial filters of *W* (first column vectors) maximizing the signal variance for group 1 and minimizing the signal variance for group 2, and the rightmost spatial filters (last column vectors) maximizing the signal variance for group 2 and minimizing the signal variance for group 1. For the CSP analyses depicted on Figure 4, each participant’s final score was calculated as the average CSP power of their epochs.

### Linear classifier

A Support Vector Machine with a linear kernel (SVM-L) was the classifier of choice throughout this work. SVMs are useful for data classification as they find the separating hyperplane with the maximal margin between two classes of data. Given a set of data X_*i*_ with corresponding labels y_i_ ∈ {1, −1}, SVM solves the unconstrained problem:

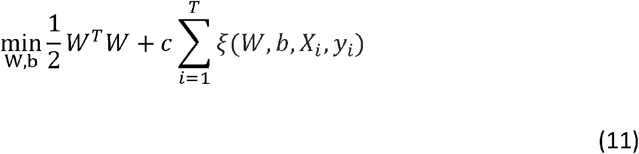

where *T* is the number of epochs, ξ(W, b, X_*i*_, y_*i*_) is a loss function and c ≥ 0 is a regularization parameter on the training error. The loss function used in this work was a L2-loss:

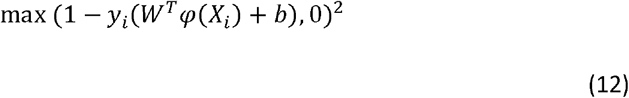

where φ is the function that maps the training data into a higher dimensional space in non-linear instances of SVM. In the linear case, however, *φ*(*X_i_*) = *X_i_*. Therefore, for any testing instance *x*, the predictor function for SVM-L is similar to that used in linear discriminant analysis:

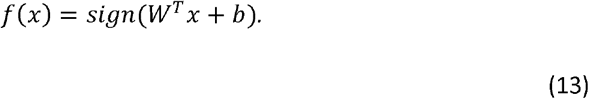

To optimize the model training against the class imbalances present on our datasets (especially the story-listening task), the regularization parameter was balanced for each class. Each class weight would take the proportion of class frequencies into account and the new parameter *c’* is then calculated as *c*’ = *class_weight*[*i*] * *c*. In practice, the SVM-L for classification was applied using Python’s *scikit-learn* package implementation (*svm.SVC* function).

Grid search cross-validation was used to optimize not only the regularisation parameter but also the number of features to use for the CSP (i.e., filters) as only a subset of the total filters is used (viz. the *m* first and last rows of S, i.e., S_p_, p ∈{1…2*m*}). This cross-validation process was denoted as Leave-One-Subject-Out cross-validation since, for each fold, all epochs of a single participant were held for validation while the other epochs were used for training the classifier. The label attributed to the child (e.g., dyslexic / non-dyslexic) was based on the majority of classifications for that child’s epochs. For every child, the proportion of story listening epochs classified as “dyslexic” was used to create the receiver operating characteristic (ROC) curves. The feature estimation process was also cross-validated between 1) variance, 2) log variance and 3) proportional variances as in [91]:

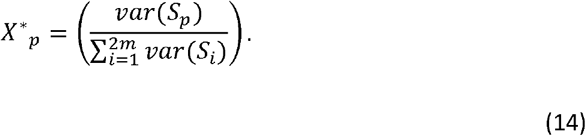

For both datasets, the variance features worked best. For all classifiers, the other hyperparameters were kept at *m* = 2, *c* = 100 as determined by grid-search cross-validation. During this feature engineering process we also found that normalizing variables by a power of 10 helped model convergence in some cases. This was the case for the story listening task, where this constant was set to 10e4 after grid-search crossvalidation.

## Data and code accessibility

All original data and code will be deposited and made publicly available at the date of publication.

Any additional information required to reanalyze the data reported in this paper is available from the lead contact upon request.

## Acknowledgements

The research was funded by a donation from the Yidan Prize Foundation to U.G. Collection of the EEG datasets was funded by a grant awarded to U.G. by the Fondation Botnar (project number 6064) and an Australian Research Council Discovery Project grant (DP110105123) awarded to U.G. and D.B. B.D.S is funded by a Royal Society E.P. Abraham Research Professorship (RP/R1/180165). M.K. receives support from the Basque Government through the BERC 2018-2021 program, the Spanish State Research Agency through BCBL Severo Ochoa excellence accreditation CEX2020-001010-S, and the Spanish Ministry of Science and Innovation through the Ramon y Cajal Research Fellowship, RYC2018-024284-I. G.D.L. work was conducted with the financial support of Science Foundation Ireland under Grant Agreement No. 13/RC/2106_P2 at the ADAPT SFI Research Centre at Trinity College, The University of Dublin. ADAPT, the SFI Research Centre for AI-Driven Digital Content Technology, is funded by Science Foundation Ireland through the SFI Research Centres Programme. The sponsors played no role in the study design, data interpretation or writing of the report. The authors would like to thank all the children and families involved in the study.

## Notes

### Competing Interest Statement

The authors have declared no competing interest.

