## Supplementary material for "Atypical cortical encoding of speech identifies children with Dyslexia versus Developmental Language Disorder": Table S1

**Table S1** – Description of the full sample of children included in the story listening task analyses. The table shows the individual data for subgroups of typically-developing (CTR), dyslexic (DYS) and DLD children (DLD).

| Group | Child Code | Age (months) | PH_AW[a] | TOWRE[b] | | TROG[c] | *WIAT*-VOCAB[d] | CELF *SENTENCES*[e] | NVIQ[f] |
|---|---|---|---|---|---|---|---|---|---|
| | | | | *W* | *NW* | | | | |
| CTR (RL) | CTR-01 | 92.7 | 90 | 107 | 101 | 111 | 120* | 14 | 115 |
| CTR (RL) | CTR-02 | 90.6 | 94 | 91 | 93 | 106 | 100* | 10 | 122 |
| CTR (CA) | CTR-03 | 143.2 | 103 | 122 | 111 | 106 | 90 | 8 | 108 |
| CTR (RL) | CTR-04 | 76.6 | 110 | 90 | 92 | 98 | 123 | 13 | 85 |
| CTR (RL) | CTR-05 | 86.4 | 103 | 90 | 93 | 125 | 109 | 8 | 131 |
| CTR (CA) | CTR-06 | 96.2 | 90 | 103 | 87 | 99 | 107 | 10 | 116 |
| CTR (CA) | CTR-07 | 124.2 | 107 | 100 | 101 | 116 | 119 | 14 | 111 |
| CTR (RL) | CTR-08 | 86.9 | 103 | 120 | 118 | 102 | 88 | 9 | 99 |
| CTR (RL) | CTR-09 | 80.7 | 112 | 138 | 121 | 109 | 113 | 14 | 113 |
| CTR (RL) | CTR-10 | 81.3 | 110 | 112 | 80 | 99 | 113 | 11 | 109 |
| CTR (CA) | CTR-11 | 104.2 | 94 | 89 | 90 | 109 | 88 | 8 | 100 |
| CTR (RL) | CTR-12 | 72.7 | 105 | 87 | 92 | 111 | 107 | 8 | 102 |
| CTR (CA) | CTR-13 | 120.2 | 103 | 110 | 111 | 92 | 107 | 9 | 108 |
| CTR (CA) | CTR-14 | 105.2 | 127 | 126 | 124 | 113 | 131 | 15 | 124 |
| CTR (CA) | CTR-15 | 111.8 | 98 | 103 | 106 | 109 | 100 | 11 | 118 |
| CTR (RL) | CTR-16 | 82.2 | 129 | 100 | 95 | 109 | 92 | 11 | 109 |
| CTR (CA) | CTR-18 | 102.2 | 90 | 109 | 100 | 109 | 97 | 10 | 124 |
| CTR (CA) | CTR-18 | 118.7 | 116 | 88 | 91 | 113 | 123 | 14 | 114 |
| CTR (CA) | CTR-20 | 98.8 | 116 | 109 | 102 | 113 | 114 | 15 | 103 |
| CTR (CA) | CTR-21 | 94.8 | 105 | 112 | 103 | 111 | 97 | 10 | 124 |
| CTR (CA) | CTR-22 | 126.0 | 120 | 95 | 93 | 106 | 98 | 12 | 127 |
| CTR (CA) | CTR-23 | 136.2 | 96 | 80 | 76 | 97 | 89 | 11 | 108 |
| CTR (CA) | CTR-24 | 108.0 | 116 | 85 | 94 | 116 | 102 | 16 | 124 |
| CTR (CA) | CTR-25 | 103.7 | 112 | 91 | 100 | 99 | 104 | 11 | 105 |
| CTR (RL) | CTR-26 | 85.8 | 107 | 79 | 93 | 106 | 95 | 7 | 105 |
| CTR (RL) | CTR-27 | 84.8 | 112 | 114 | 107 | 106 | 109 | 15 | 118 |
| CTR (CA) | CTR-28 | 96.6 | 100 | 96 | 96 | 109 | 100 | 10 | 119 |
| CTR (CA) | CTR-29 | 116.6 | 84 | 92 | 81 | 118 | 102 | 11 | 98 |
| CTR (RL) | CTR-30 | 76.3 | 110 | 101 | 104 | 120 | 102 | 11 | 103 |
| CTR (RL) | CTR-31 | 85.6 | 96 | 85 | 90 | 102 | 102 | 11 | 112 |
| CTR (CA) | CTR-32 | 111.2 | 100 | 94 | 101 | 104 | 107 | 11 | 118 |
| CTR (CA) | CTR-33 | 128.0 | 110 | 85 | 87 | 111 | 110 | 17 | 114 |
| CTR (CA) | CTR-35 | 105.4 | 110 | 108 | 106 | 85 | 102 | 13 | 100 |
| CTR (CA) | CTR-36 | 98.7 | 103 | 120 | 111 | 109 | 114 | 12 | 110 |
| CTR (CA) | CTR-37 | 125.0 | 94 | 118 | 116 | 92 | 112 | 12 | 111 |
| CTR (CA) | CTR-38 | 116.9 | 90 | 103 | 113 | 109 | 131 | 10 | 114 |
| CTR (CA) | CTR-39 | 130.1 | 100 | 138 | 131 | 92 | 117 | 11 | 126 |
| CTR (CA) | CTR-40 | 110.3 | 100 | 91 | 121 | 113 | 107 | 13 | 129 |

| | | | | | | | | | |
|---|---|---|---|---|---|---|---|---|---|
| CTR (RL) | CTR-01 | 89.9 | 114 | 108 | 99 | 106 | 109 | 12 | 91 |
| CTR (CA) | CTR-01 | 121.8 | 112 | 100 | 113 | 106 | 107 | 12 | 121 |
| DYS | DYS-01 | 98.7 | 75 | 65 | 73 | 85 | 90* | 4 | 103 |
| DYS | DYS-02 | 110.5 | 86 | 68 | 75 | 113 | 105* | 12 | 95 |
| DYS | DYS-03 | 137.5 | 90 | 80 | 73 | 102 | 90 | 6 | 115 |
| DYS | DYS-04 | 103.9 | 94 | 91 | 73 | 90 | 100* | 9 | 111 |
| DYS | DYS-05 | 120.6 | 88 | 61 | 65 | 106 | 129 | 8 | 110 |
| DYS | DYS-06 | 126.8 | 96 | 65 | 81 | 116 | 112 | 12 | 103 |
| DYS | DYS-07 | 120.8 | 114 | 63 | 78 | 106 | 107 | 13 | 112 |
| DYS | DYS-08 | 86.3 | 86 | 76 | 74 | 97 | 102 | 8 | 107 |
| DYS | DYS-09 | 115.5 | 82 | 75 | 66 | 90 | 111 | 11 | 91 |
| DYS | DYS-10 | 124.7 | 98 | 80 | 83 | 111 | 92 | 9 | 124 |
| DYS | DYS-11 | 97.4 | 75 | 93 | 84 | 104 | 93 | 12 | 100 |
| DYS | DYS-12 | 119.3 | 88 | 90 | 82 | 104 | 102 | 12 | 107 |
| DYS | DYS-13 | 129.4 | 77 | 81 | 83 | 111 | 110 | 13 | 114 |
| DYS | DYS-14 | 141.2 | 80 | 92 | 99 | 97 | 103 | 9 | 99 |
| DYS | DYS-15 | 120.3 | 65 | 70 | 78 | 88 | 86 | 7 | 94 |
| DYS | DYS-16 | 72.0 | 82 | 87 | 84 | 97 | 107 | 6 | 93 |
| DLD | DLD-01 | 123.7 | 92 | 100 | 107 | 97 | 107 | 5 | 110 |
| DLD | DLD-02 | 84.8 | 88 | 100 | 110 | 69 | 95 | 6 | 91 |
| DLD | DLD-03 | 101.7 | 100 | 98 | 101 | 85 | 84 | 7 | 97 |
| DLD | DLD-04 | 95.3 | 84 | 81 | 81 | 69 | 105* | 6 | 91 |
| DLD | DLD-05 | 81.6 | 105 | 90 | 100 | 109 | 85 | 6 | 118 |
| DLD | DLD-06 | 98.1 | 90 | 97 | 84 | 67 | 75 | 10 | 83 |
| DLD | DLD-07 | 79.5 | 84 | 95 | 80 | 76 | 80 | 11 | 103 |

RL – Reading Level; CA – Chronological Age

[a] Comprehensive Test of Phonological Processing (M = 100, SD = 15)

[b] Test of Word Reading Efficiency – word (*W*) and non-word (*NW*) reading (M = 100, SD = 15)

[c] Test of Receptive Oral Grammar (M = 100, SD = 15)

[d] WIAT Vocabulary (M = 100, SD = 15)

[e] Clinical Evaluation of Language Fundamentals, sentence recall (M = 10, SD = 3)

[f] Kaufman Brief Intelligence Test (M = 100, SD = 15)

*****Interpolated scores, as these children were tested with a different vocabulary instrument with a mean of 10, SD 3. To match the standardization of the WIAT, which has a mean of 100 and SD 15, a score of 10 is converted to 100, a score of 11 to 105, and so on.
