## Supplementary material for "Atypical cortical encoding of speech identifies children with Dyslexia versus Developmental Language Disorder": Table S2

**Table S2** – Description of the full sample of children included in the rhythmic syllable task analyses. The table shows the individual data for subgroups of typically-developing (CTR) and dyslexic (DYS) children.

| Group | Child Code | Age (months) | PHAB_R[a] | FSIQ[b] | BPVS[c] | BAS[d] | | TOWRE[e] | |
|---|---|---|---|---|---|---|---|---|---|
| | | | | | | *R* | *S* | *W* | *NW* |
| CTR | CTR-1 | 108.6 | 107 | 118.85 | 121 | 113 | 109 | 110 | 107 |
| CTR | CTR-2 | 109.8 | 102 | 96.31 | 95 | 99 | 92 | 106 | 111 |
| CTR | CTR-3 | 106.6 | 102 | 86.65 | 81 | 97 | 108 | 109 | 106 |
| CTR | CTR-4 | 120.1 | 108 | 110.8 | 104 | 97 | 89 | 94 | 96 |
| CTR | CTR-5 | 108.2 | 83 | 97.92 | 110 | 106 | 96 | 101 | 112 |
| CTR | CTR-6 | 109.1 | 101 | 105.97 | 100 | 101 | 95 | 109 | 93 |
| CTR | CTR-7 | 113.5 | 110 | 104.36 | 109 | 97 | 96 | 95 | 97 |
| CTR | CTR-8 | 110.5 | 101 | 101.14 | 121 | 114 | 90 | 114 | 107 |
| CTR | CTR-9 | 110.7 | 102 | 97.92 | 102 | 97 | 96 | 94 | 93 |
| CTR | CTR-10 | 114.2 | 104 | 109.19 | 116 | 101 | 89 | 101 | 89 |
| CTR | CTR-11 | 116.6 | 99 | 85 | 83 | 88 | 88 | 92 | 97 |
| CTR | CTR-12 | 111.3 | 107 | 97.92 | 96 | 103 | 96 | 105 | 109 |
| CTR | CTR-13 | 110.6 | 107 | 125.29 | 113 | 91 | 100 | 102 | 86 |
| CTR | CTR-14 | 109.0 | 106 | 114.02 | 102 | 96 | 94 | 91 | 103 |
| CTR | CTR-15 | 112.8 | 94 | 102.75 | 98 | 96 | 103 | 88 | 90 |
| CTR | CTR-16 | 95.3 | 101 | 110.8 | 100 | 97 | 97 | 111 | 100 |
| CTR | CTR-17 | 106.7 | 103 | 105.97 | 97 | 95 | 102 | 90 | 88 |
| CTR | CTR-18 | 101.9 | 104 | 114.02 | 102 | 101 | 94 | 105 | 85 |
| CTR | CTR-19 | 102.2 | 106 | 85.04 | 98 | 95 | 99 | 97 | 92 |
| CTR | CTR-20 | 110.1 | 102 | 101.14 | 121 | 102 | 98 | 105 | 105 |
| CTR | CTR-21 | 103.8 | 106 | 115.63 | 100 | 103 | 107 | 104 | 92 |
| DYS | DYS-1 | 112.2 | 101 | 105.97 | 84 | 85 | 78 | 91 | 81 |
| DYS | DYS-2 | 98.6 | 80 | 97.92 | 78 | 73 | 67 | 66 | 55 |
| DYS | DYS-3 | 113.6 | 103 | 110.8 | 117 | 86 | 81 | 74 | 81 |
| DYS | DYS-4 | 111.8 | 103 | 109.19 | 115 | 82 | 78 | 76 | 63 |
| DYS | DYS-5 | 102.6 | 79 | 101.14 | 99 | 79 | 86 | 77 | 96 |
| DYS | DYS-6 | 116.1 | 94 | 86.65 | 117 | 84 | 79 | 77 | 82 |
| DYS | DYS-7 | 101 | 106 | 94.7 | 88 | 89 | 82 | 99 | 76 |
| DYS | DYS-8 | 117 | 78 | 94.7 | 110 | 70 | 72 | 68 | 73 |
| DYS | DYS-9 | 99 | 84 | 89.87 | 102 | 96 | 90 | 105 | 71 |
| DYS | DYS-10 | 99.1 | 84 | 91.48 | 97 | 83 | 84 | 94 | 88 |
| DYS | DYS-11 | 116 | 104 | 105.97 | 118 | 81 | 78 | 88 | 70 |
| DYS | DYS-12 | 111.6 | 89 | 112.41 | 87 | 67 | 75 | 55 | 55 |
| DYS | DYS-13 | 116 | 78 | 115.63 | 97 | 74 | 73 | 71 | 87 |
| DYS | DYS-14 | 113 | 101 | 114.02 | 101 | 88 | 75 | 82 | 81 |
| DYS | DYS-15 | 117.9 | 101 | 105.97 | 115 | 83 | 73 | 92 | 86 |
| DYS | DYS-16 | 116.9 | 70 | 93.09 | 104 | 80 | 83 | 79 | 81 |
| DYS | DYS-17 | 107.6 | 81 | 96.31 | 102 | 91 | 84 | 90 | 82 |
| DYS | DYS-18 | 107.4 | 79 | 94.7 | 108 | 77 | 74 | 65 | 71 |
| DYS | DYS-19 | 110.3 | 99 | 94.7 | 102 | 70 | 74 | 60 | 75 |
| DYS | DYS-20 | 106.5 | 73 | 96.31 | 94 | 75 | 82 | 71 | 74 |
| DYS | DYS-21 | 105.6 | 81 | 97.92 | 113 | 76 | 77 | 95 | 76 |

| | | | | | | | | | |
|---|---|---|---|---|---|---|---|---|---|
| DYS | DYS-22 | 95 | 102 | 110.8 | 112 | 89 | 84 | 79 | 84 |
| DYS | DYS-23 | 121.1 | 94 | 85.04 | 102 | 67 | 65 | 55 | 71 |
| DYS | DYS-24 | 107 | 102 | 107.58 | 96 | 77 | 81 | 75 | 77 |
| DYS | DYS-25 | 110 | 99 | 107.58 | 121 | 86 | 90 | 88 | 79 |
| DYS | DYS-26 | 108.5 | 101 | 97.92 | 101 | 79 | 78 | 95 | 86 |
| DYS | DYS-27 | 110.7 | 81 | 126.9 | 121 | 82 | 78 | 78 | 76 |

[a] Phonological Assessment Battery: Rhyme Awareness
[b] Full Scale IQ
[c] British Picture Vocabulary Scale
[d] British Ability Scales: Reading (*R*) and Spelling (*S*)
[e] Test of Word Reading Efficiency: Words (*W*) and Non-Words (*NW*)
For all tests M = 100; SD = 15.
